# An open-source stereotaxic container with an integrated cutting guide for human brain fixation during magnetic resonance imaging and sectioning for histology

**DOI:** 10.64898/2026.06.01.729391

**Authors:** Jacob Berardinelli, Jr-Jiun Liou, Nadim Farhat, Jinghang Li, Mark Stauffer, Karla L. Miller, Benjamin C. Tendler, Wenchuan Wu, Milos D. Ikonomovic, Joseph M. Mettenburg, Howard J. Aizenstein, Tales Santini, Julia K. Kofler, Tamer S. Ibrahim

## Abstract

**Introduction:** Postmortem imaging at ultrahigh field strengths, such as 7 Tesla (7T) magnetic resonance imaging (MRI), enables unprecedented visualization of fine brain structures and pathology. However, precise registration between MRI and histology is often compromised by tissue deformation and lack of standardized tools for preserving anatomical orientation. A reliable, high-resolution MRI-to-histology workflow is essential for MRI-guided sampling to investigate neurodegenerative diseases.

**Methods:** We developed a human postmortem brain container composed of stereotaxic enclosure and cutting guide system using iterative 3D printing. The final design is constructed from Victrex AM™ 200, a polyaryletherketone material engineered with MRI compatibility, high mechanical strength, and high resistance to fixatives. Human brains were embedded in an agar/sucrose mixture within the container and scanned at 7T using high-contrast and high-resolution imaging sequences. The cutting guide enables reproducible sectioning at defined tissue coronal slab thicknesses (9.2 ± 0.6 mm). Block face photographs of each coronal slab were then co-registered with MRI for targeted tissue sampling.

**Results:** To date, 215 human postmortem brains have been imaged ex vivo using this container. The container maintains consistent spatial orientation from imaging through sectioning, supporting correlation between MR-visible features and histological findings. Imaging-visible abnormalities, such as white matter hyperintensities and cerebral microbleeds, were localized and sampled with minimal deformation. The container enables single-scan workflows, avoiding custom cutting templates or repeat imaging. It supports deep learning-based lesion segmentation and MRI-guided histology registration, facilitating large-scale neuropathological investigations.

**Conclusion:** The stereotaxic cutting guide system addresses a critical gap in postmortem imaging by integrating MRI-compatible materials, precision-guided sectioning, and high-throughput imaging. It has enabled multimodal studies of aging and neurodegeneration and is being adopted in neuropathological workflows. This platform provides a reproducible, scalable, and anatomically precise solution for aligning ultrahigh field MRI with histology, enhancing both clinical research and digital pathology efforts.

## Introduction

Postmortem magnetic resonance imaging (MRI) serves as an intermediate modality between in vivo neuroimaging and postmortem histopathology, providing high-resolution anatomical imaging that maintains spatial correspondence with tissues for microscopic examination [1–3]. This intermediary role enables postmortem MRI to validate in vivo imaging findings, when available, and guide targeted histological sampling of regions invisible to gross anatomical inspection [4, 5]. Moreover, postmortem MRI provides several technical advantages over in vivo imaging, including the ability to scan at ultra-high resolutions due to longer imaging times, and the absence of motion artifacts [2, 6]. In addition, 7 Tesla (7T) MRI is a powerful tool for postmortem imaging of the brain as it provides improved image resolution and contrast [2, 7]. Still, postmortem imaging faces unique challenges differentiating it from in vivo tissue; the postmortem brain tissue can experience dehydration, membrane breakdown, axonal beading, compaction of extracellular space, shrinkage, and increased rigidity as a result of the postmortem interval and chemical fixation methods [8–12].

Interpretation of findings in postmortem MRI images require accurate registration with histology sections, which provide information about the underlying cellular and molecular changes. Registration of postmortem MRI images to histology sections is a complex process [13–21] and is critical for several applications, including the study of neurodegenerative diseases, traumatic brain injury, and developmental disorders [17, 22–25]. Current state-of-the-art methods [16, 19, 26–28] for MRI-to-gross-anatomy registration often require multiple scanning sessions, custom anatomical conforming cutting guides, and complex histological alignment processes. Alternative postmortem brain imaging workflows image the brain floating freely in a fluid like fomblin or formalin, causing susceptibility to motion artifacts, or fix the brain with a physical mechanism which can lead to additional tissue deformations [16, 19, 26–28]. Histological registration is then done based on block face photography, leveraging deformable image registration with varying degrees of manual or automation techniques [14, 15, 29]. Moreover, some groups developed brain specific cutting guides, designed after an initial brain imaging and surface reconstruction to preserve spatial correspondence between the imaging and gross anatomy slices [17, 20, 24, 28, 30], however, this causes design and manufacturing delays between imaging and sampling.

This study leverages the development of a reusable container with cutting guides as well as embedding protocol and embedding media that allows for high quality postmortem imaging at 7T, facilitating the histopathology alignment, and is also compatible with other MRI field strength. The containers were developed to be used with regular human head coils used for in-vivo scans. To date, the containers were used in 215 ex-vivo brain scans in collaboration with the Alzheimer’s Disease Research Center (ADRC) in Pittsburgh [14, 31–39].

## Methods

### Design requirements

A central requirement for high-resolution postmortem imaging and histological registration is maintaining anatomical fidelity across multiple modalities while enabling practical, reproducible workflows. The development of the container system and integrated cutting guide was led by a series of stringent project needs: (1) compatibility with ultra-high field MRI equipment, such as the limited space within head coils, (2) provision of support for stereotaxic alignment and stability throughout the processes of agar embedding and imaging, (3) facilitation of precise and consistent tissue sectioning which supports subsequent MRI-to-histology registration, (4) extended lifespan allowing for sectioning across multiple acquisitions, and (5) facilitation of easy cleaning and disinfection procedures. The design evolved over four versions of container through an iterative prototyping process to meet these objectives.

### Container Version 1

The first functional prototype (V1) consisted of a sharp geometric lid, base, and a single cutting guide **(Fig 1)** manufactured in Acrylonitrile Butadiene Styrene (ABS). The height (right-left direction inside the coil) of the base was 80 mm, and the lid with a custom glued rubber gasket extended an additional 16 mm. The total width (anterior-posterior or AP direction inside the coil) and length (superior-inferior or SI direction inside the coil) of the container measured 200 mm and 153 mm, respectively. The initial cylindrical columns were 3 mm in diameter by 73 mm long with a 0.55 mm gap in between.

**Figure 1.**
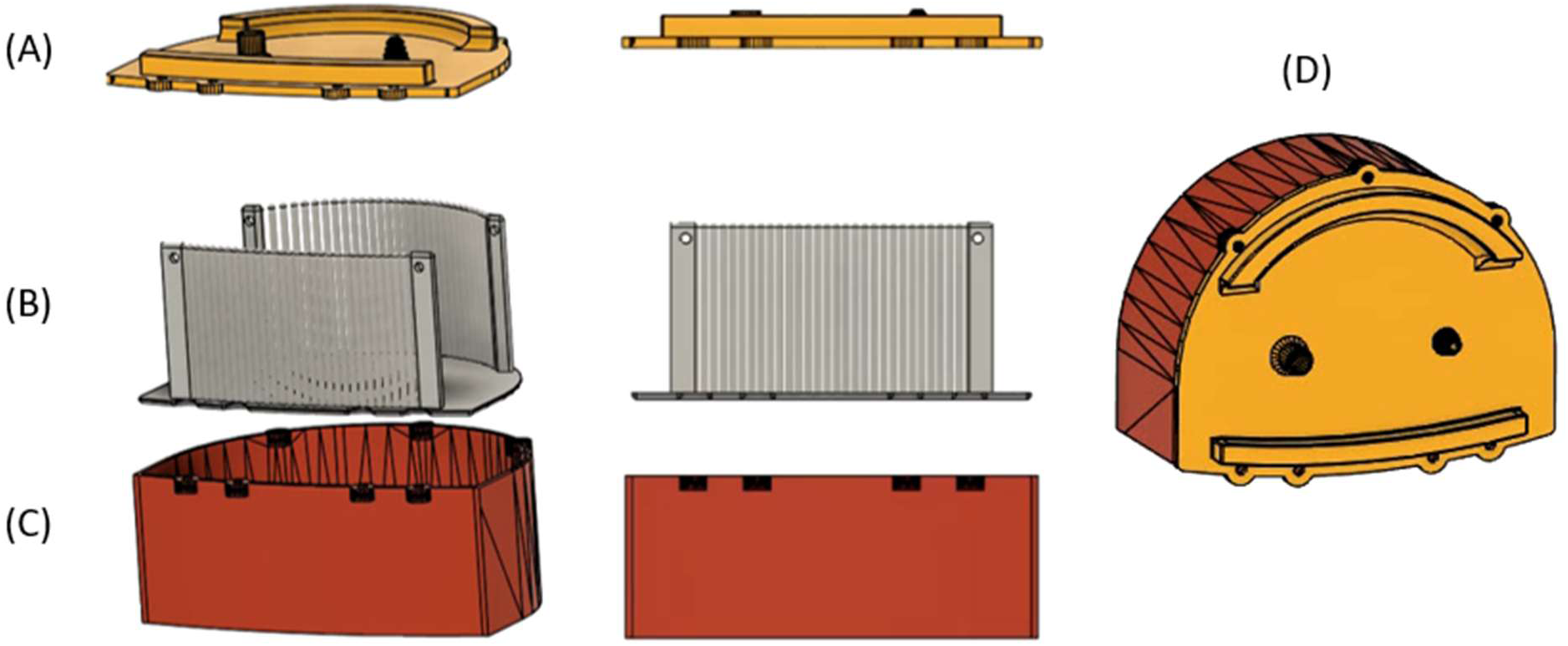
Container Version 1. The first functional postmortem container comprised of a lid (A), a cutting guide (B), and a base (C). The larger non-cylindrical columns on the anterior and posterior of the cutting guide features a hole to insert stainless steel rods for cutting guide removal from the base when filled with agar and brain tissue (D).

The 3D printed lid contained a vacuum fill port and hose barb adapter, designed for using vacuum pressure to eliminate air bubbles.

Another important feature was the implementation of non-cylindrical, stadium prism shaped columns at the anterior and posterior of the guide. These served a dual function: structurally, they supported the knife at the furthest extremes in the anterior-posterior direction, and additionally, their design allowed insertion of extraction rods from the bottom row of columns to the top row (superior-inferior direction), to facilitate removal of the cutting guide as agar adheres the guide to the base. All elongated columns protrude into the lid’s raised sections providing an air pocket that leaving them exposed above the level of the agar; thus accessible for knife alignment for all subsequent cutting.

### Dual Cutting Guide Version 1

The initial dual cutting guide addon **(Fig 2)** incorporated both coronal and axial sectioning options resulting in columns distributed nearly 360 degrees around the brain. The columns were shorter, measuring 66.5 mm and ending at the height of the base container. Only the four larger posts to remove the cutting guide from the base protruded into the lid, free from being submerged in agar. Two axial cuts were made, to remove a large brain slab for gross anatomy imaging, prior to returning it for further coronal sectioning.

**Figure 2.**
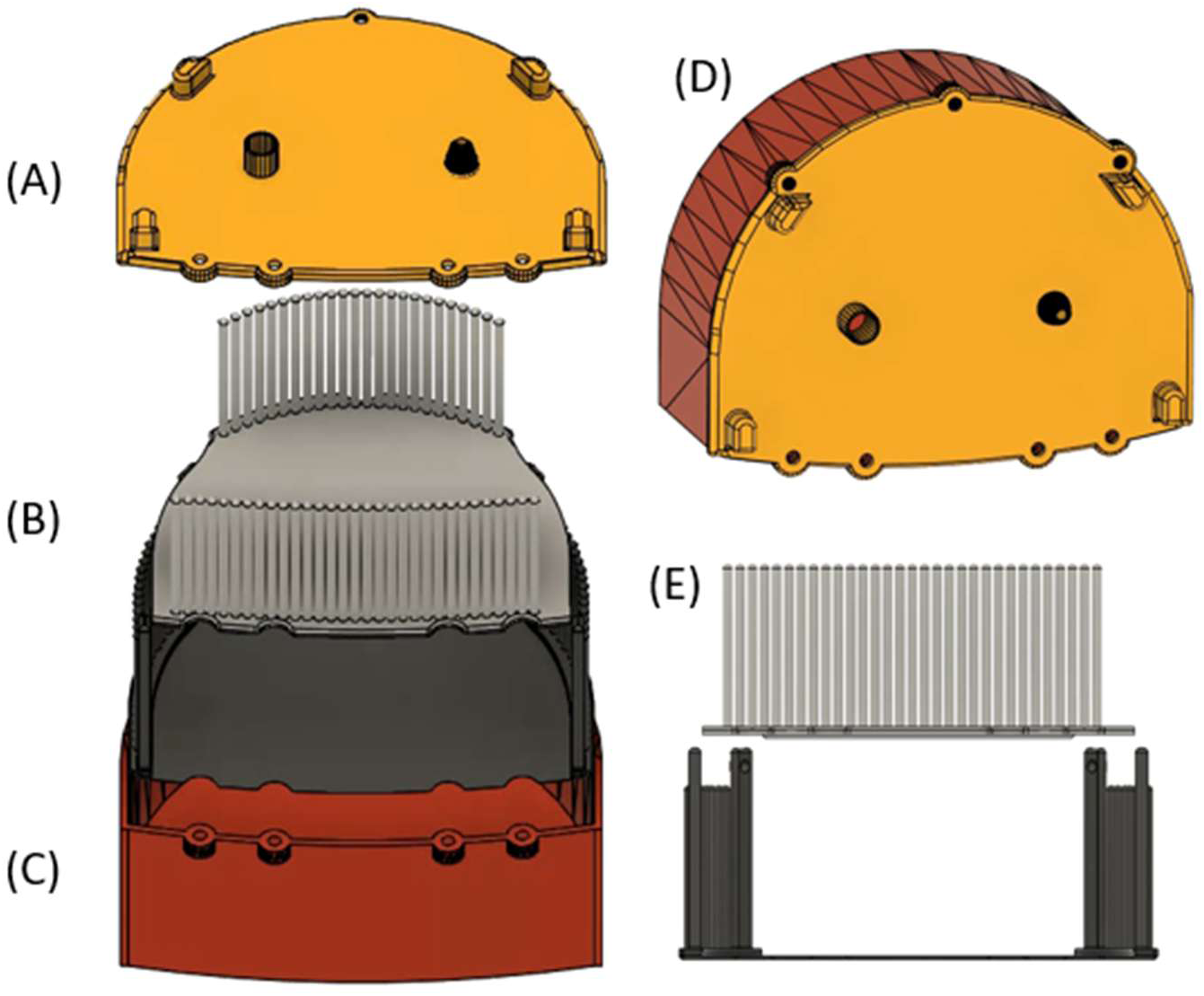
Dual Cutting Guide Version 1. It is comprised of a lid (A), dual cutting guides (B&E), and a base (C). Four longer posts featuring a hole to insert a rod for cutting guide removal are on the bottom cutting guide in the anterior-posterior direction (D).

### Container Version 2

Container Version 2 (V2) marked a transition to full polycarbonate (PC) construction **(Fig 3)**. Columns increased to 4 mm diameter and 83 mm length, and the gap between columns increased to 0.6 mm to accommodate the knife with improved tolerances without sacrificing knife rigid alignment between the columns. The height of the base decreased from 80 mm to 75 mm while integrating a domed lid design. Rounding all corners, reducing base height, and adding a domed lid increases the internal volume of the container the brain can occupy while decreasing the largest external dimensions that limited the container fitment in the rigid receive coil. A custom cut rubber gasket was glued to the inside ridge of the lid to provide a better seal between the lid and the base. The small fill port and vacuum barbed hose adapter were replaced by a larger hole covered with a screw cap.

**Figure 3.**
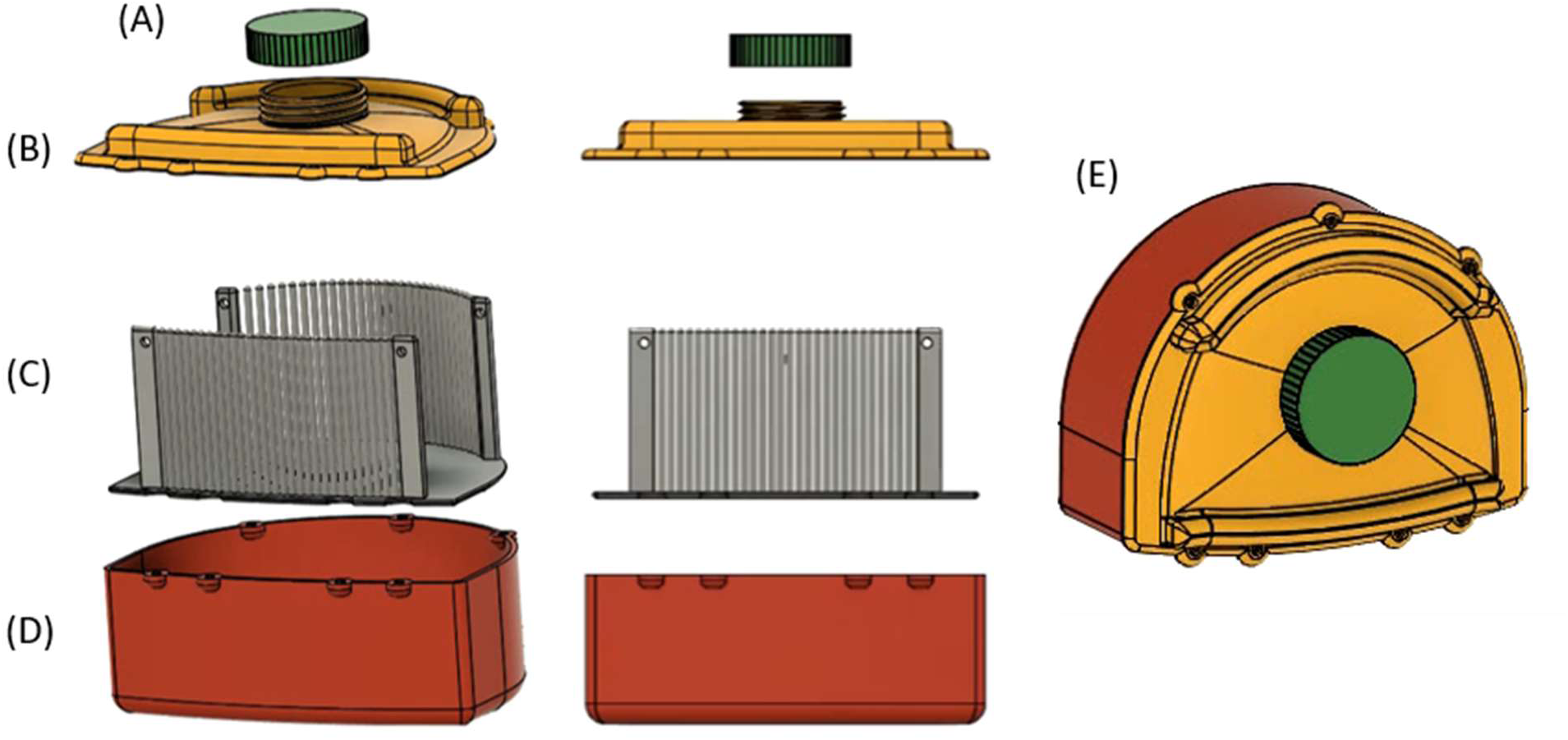
Container Version 2. It is comprised of a screw cap (A), a domed lid with rounded edges (B), cutting guides (C), and a base (D) minimizing air inside the container near brain anatomy. The cutting guide remains unchanged from Container Version 1 (E).

### Dual Cutting Guide Version 2

The V2 with the dual cutting guides **(Fig 4)** increased column length to 76.5 mm, allowing the columns to protrude above the agar surface and facilitate proper knife alignment before beginning to cut into agar. Additionally, the domed design was integrated into the dual cutting guide lid increasing internal volume and rounding of all sharp corners.

**Figure 4.**
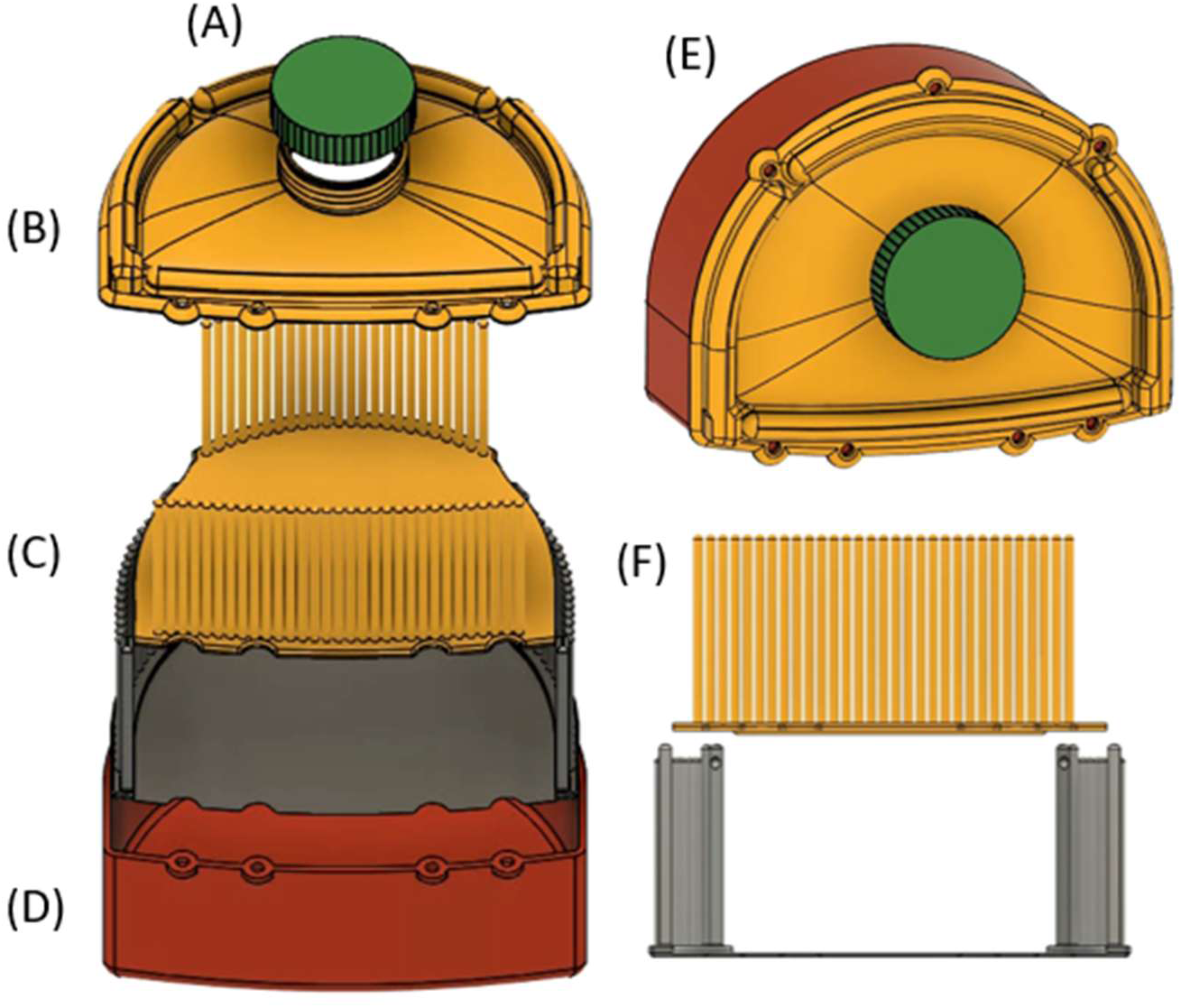
Dual Cutting Guide Version 2. It is (E) comprised of a screw cap (A), a domed lid with rounded edges (B), longer posts on both cutting guides (C&F), and a base (D).

### Container Version 3

Container Version 3 (V3) deviated significantly from previous designs to drastically reduce external volume, eliminate the need for hardware, and eliminate the custom-cut rubber gasket **(Fig 5)**. This version introduced a two-part snap-fit joint lid, entirely removing all metal hardware and threaded inserts, which had previously contributed to disruption of the radio frequency (RF) field homogeneity. The outer tabs of the first lid are bent inwards, applying pressure on the anterior and posterior walls of the base to stay in place. The second lid articulates on ridges on the side of the raised portions of the lid that the cutting guide columns insert into.

**Figure 5.**
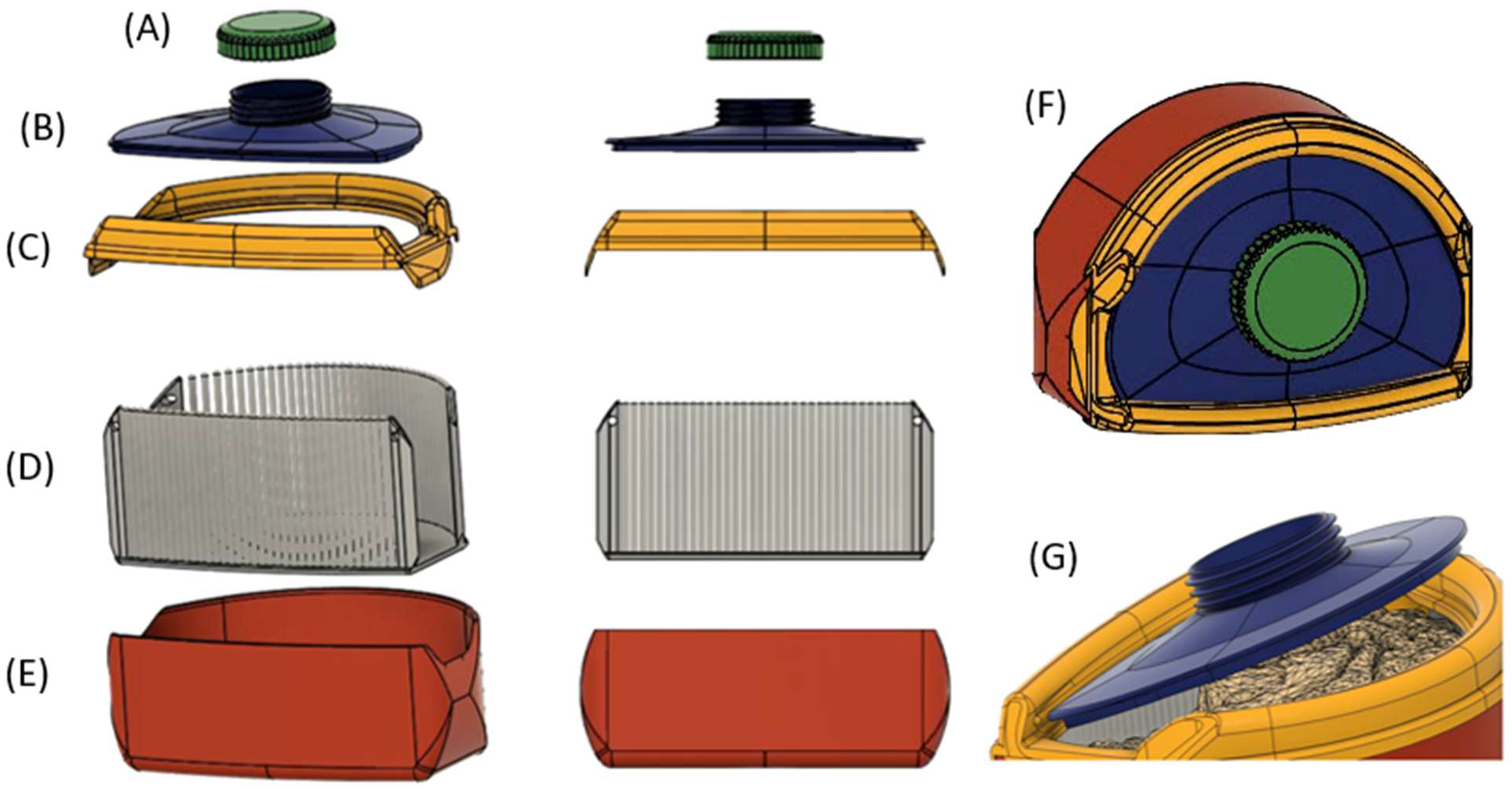
Container Version 3. It comprised of a screw cap (A), a two piece snap fit lid (B, C, G), eliminating hardware and adding 6 columns on each row of the cutting guide (D) while decreasing external volume by further rounding the base (E) lid and screw cap while slanting the larger posts with holes for cutting guide removal (F).

Once pressed into place the smaller ridge on top holds the lid in a snap fit (Fig 5G & Supp Fig 1).

To enable removal of the cutting guide with an ellipsoid base and lid shape, cutouts were added to the anterior and posterior walls of the base, allowing the guide, agar, and brain to easily slide out while the lid replaced the missing walls. These cutouts initially only allowed the base to be partially filled and subsequently the brain only partially submerged. That is until the first lid (Fig 5C) is snapped into place, as the brain floating in agar needs to be held, submerged, as the agar continues to fill the container to the opening of the first lid until the agar solidifies. It is not until the brain no longer floats that the second lid piece (Fig 5B) can be snaped into place and final agar added before the screw cap (Fig 5A) seals the container closed. The number of columns increased from 28 per row in V2 to 34 per row in V3, providing enhanced spatial precision during sectioning. The outer posts, used for extracting the cutting guide with rods, were significantly enlarged to facilitate rod insertion through the bottom row of columns, and these posts were slanted to prevent the lid from protruding excessively on the corners, moving further from a rectangular prism to ellipsoid.

### Container Version 4 (Final)

Container Version 4 (V4) is the culmination of iterative engineering, integrating the longer 82.5 mm columns and rounded base from V3 while reverting to a fully enclosed one-piece lid by removing anterior and posterior wall cutouts **(Fig 6)**. Prototyped in PC and manufactured from Victrex AM^TM^ 200, the container achieves high mechanical strength and MRI compatibility (See Materials section). The single piece domed lid, trimmed on the anterior and posterior edges for a precise fit within the receive coil, features a 4 mm internal lip that forms a secure, gasket-free seal with the base due to stiffer plastic. Optimized thread engagement with smaller diameter, longer titanium screws minimize metallic content while ensuring structural integrity and leak resistance.

**Figure 6.**
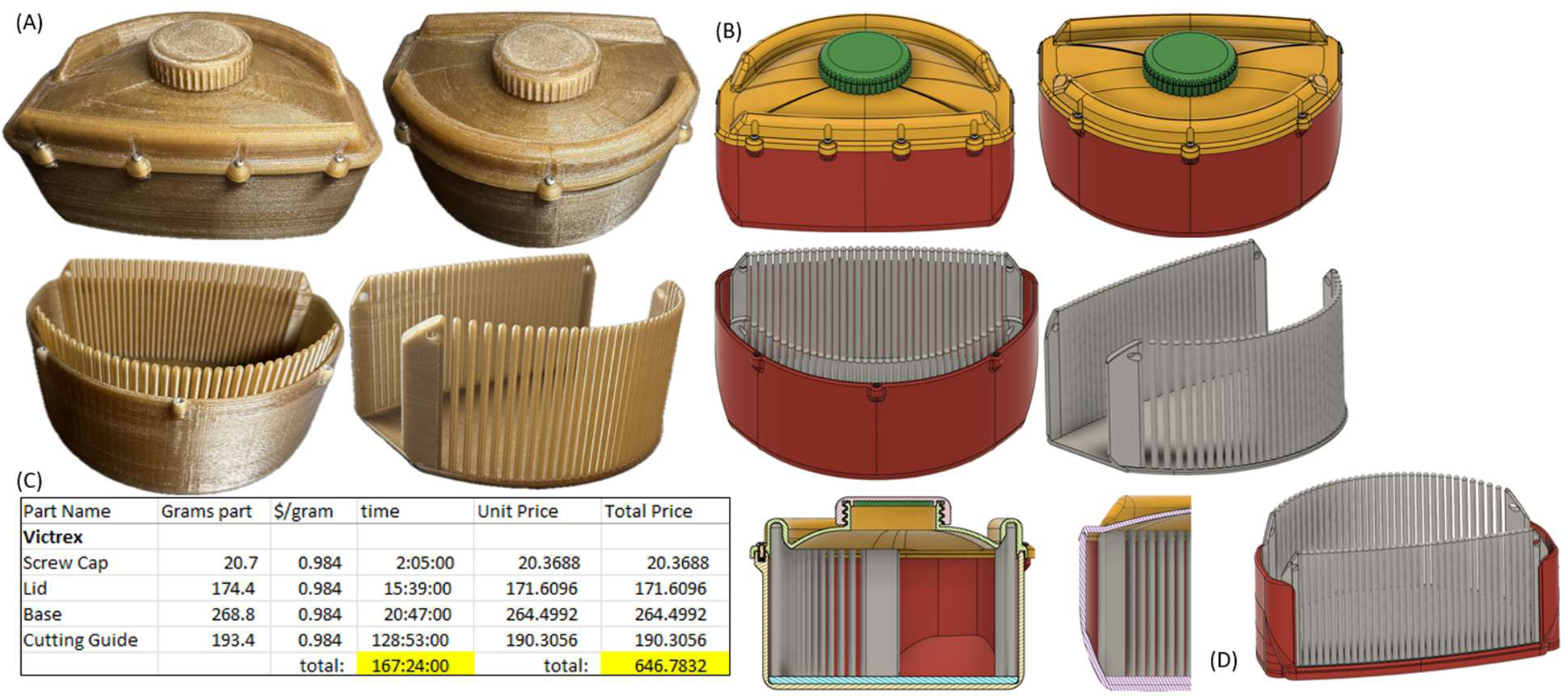
Container Version 4. (A) Images of Container Version 4 with cutting guide manufactured out of Victrex AM^TM^ 200, (B) CAD diagram of Container Version 4 design and additional post imaging cutting guide support structure (D), and (C) printing time in hours and total cost in US dollars for printing Container Version 4.

The enhanced curvature of the base facilitates container positioning further into the top of the receive towards isocenter of the transmit coil compared to V2. The slanted anterior and posterior edges of the bottom of the cutting guide conform to the enhanced curvature of the base; this has enabled easier removal after imaging with less suction than previously experienced. To prevent displacement of the overhanging agar during deep anterior–posterior sectioning, an auxiliary stabilizing insert clamps the cutting guide and agar together in the AP direction after removal of the cutting guide from the base. This tool maintains agar integrity and brain stability throughout the cutting process. Together, these design advancements produce a robust, MRI-invisible, and workflow-optimized container that enables reproducible brain embedding, high-resolution imaging, and accurate guided sectioning. An animation video demonstrating V4 (See Supp Fig 2) and another animation video comparing V2 and V4 in the receive coil (See Supp Fig 3) are included in supplementary information.

### Materials

Material considerations for 7T MRI-compatible 3D-printed containers require evaluation of mechanical properties, dielectric properties, magnetic susceptibility, and chemical resistance. ABS was initially used for ease of fused deposition modeling (FDM) manufacturing and cost to develop a proof of concept to gauge the design’s initial feasibility and operational lifetime. An upgrade from ABS, PC is commonly used in medical prototyping due to its ease of processing, moderate strength (∼60–70 MPa), and electromagnetic transparency [40, 41]. However, its relatively higher RF loss tangent can potentially lower the performance of the RF coil during ultra-high field MRI, particularly in close proximity to transmit coils [45]. PC is also susceptible to creep and environmental stress cracking leading to dimensional deformation over time under mechanical stress, thermal cycling, and degradation with repeated bleach exposure while disinfecting [40, 42].

Polyaryletherketone (PAEK) is the chemical designation of a family of high-performance aromatic polymers that includes, but is not limited to, PAEK, polyetheretherketone (PEEK), and polyetherketoneketone (PEKK) [43–46]. Victrex AM^TM^ 200 is a high performance engineered thermoplastic, designed with a lowered melting temperature without changing the glass transition temperature, designated as an LMPAEK filament developed specifically for polymer crystallization and interlayer adhesion in additive manufacturing [43, 45, 47]. Victrex AM^TM^ 200 also demonstrates improved layer adhesion and slower intrinsic crystallization leading to reduced warping compared to conventional PEEK/PAEK grades. A 300% increase in layer strength compared to Victrex’s 450G PEEK (general purpose grade PEEK) was documented in “small-scale” [45] printed features such as the thin columns of the cutting guide. The resultant isotropy of Victrex AM^TM^ 200 parts across both the z axis and xy printing plane facilitates reliable fabrication and functional longevity regardless of print or stress orientation in fused filament fabrication (FFF) [45, 48]. Although direct interlayer bonding comparisons are not available because polycarbonate and Victrex AM^TM^ 200 LMPAEK belong to entirely different classes of high-performance thermoplastics, practical experience consistently shows that long thin columns are among the most failure prone geometries in FFF when printing parameters are optimized to maintain external dimensions. Print quality, chamber conditions, and slicer optimization dominate any given part’s mechanical performance [49] for that given material. Based on anecdotal observations derived from user experience, slicer optimizing dictates that the most optimized PC print can perform no better than, and often worse than, suboptimal amorphous printing conditions for a Victrex AM^TM^ 200 part, particularly in regard to interlayer adhesion. Table 1 summarizes the material properties of ABS, PC, and PAEK.

**Table 1.** Comparison of material properties used from container and cutting guide version 1 to 4 for MR imaging and coronal tissue sectioning. Note that material properties are approximate and vary for reasons such as material brand, printer used print orientation, and model geometry.

| Material Comparison | Acrylonitrile Butadiene Styrene (ABS) | Polycarbonate (PC) | Polyaryletherketone (PAEK) |
| --- | --- | --- | --- |
| Container Version(s) | V1 | V2,V3 | V4 |
| Tensile strength | ~40 MPa | ~60-70 MPa | ~100 MPa |
| Interlayer adhesion (z) | Moderate | Moderate | High (engineered) |
| Chemical Resistance | Low-Moderate | Moderate | High |
| Sterilization | Limited | Limited | Bleach, Autoclave, |
| FFF Printability | Easy | Moderate | Complex (High temp, Crystallization optimization) |
| Material Cost | Low | Low-Moderate | High |
| ISO 10993 / USP Class VI | No | No | Yes |

In terms of MRI compatibility, Victrex AM^TM^ 200 LMPAEK belongs to the same polymer family as PEEK and PEKK, which are well documented to exhibit low magnetic susceptibility and minimal RF coupling in clinical and research MRI systems [50, 51]. Victrex AM^TM^ 200 through material testing has a dielectric constant of approximately 3.11 and loss tangent of 0.0013 [50], PAEK polymers in general have demonstrated negligible interference with imaging and no significant heating during MRI exposure [52]. Additionally, Victrex AM^TM^ 200 meets ISO 10993 and USP Class VI standards, allowing safe use in contact with biological tissue and compatibility with autoclave [47, 48]. It also shows resistance to chemical degradation in the presence of formalin, ethanol, and other fixatives routinely used in postmortem workflows [47].

While PC, polypropylene, ABS, and polymethyl methacrylate are commonly used in MRI settings, Victrex AM^TM^ 200 LMPAEK offers higher mechanical and thermal performance, superior chemical resistance, stronger interlayer adhesion, and well characterized MRI and RF compatibility, making it better suited for MR tools and integrated radiolucent components within RF coil assemblies. Its higher strength and reusability, given its compatibility with autoclaving for instance, make it advantageous for thin high duty products engineered for unrestricted reuse and sterilization. Therefore, Victrex AM^TM^ 200 LMPAEK was selected for this application due to its superior performance across the above-mentioned attributes.

### Iterative Design leveraging FFF additive manufacturing

Version 4 parts were 3D modeled using Fusion 360 (parent company, location), sliced with Intamsuite (parent company, location) NEO and printed in-house with the Intamsys Funmat Pro 410, making use of its dual extrusion capabilities. Two spools of Victrex AM^TM^ 200 were dried by heating at 125 °C for 24-72 hours before loading them into the filament chamber to ensure there is no moisture left inside the plastic before printing. The left 0.6 mm nozzle is used to print everything but the cutting guide columns. The larger nozzle diameter and layer height intrinsically result in increased tolerances for sufficient bed adhesion. Warping due to insufficient bed adhesion is a particular issue with high-performance thermoplastic polymers and all filaments in the PAEK family. Vision Miner’s nano polymer adhesive along with calibrating the first layer height, speed, extrusion multiplier/flow rate, build plate, and chamber temperatures were enough to eliminate warping but were not enough on their own for reproducible manufacturing. Printing on a ceramics glass plate with enough bed adhesion to eliminate warping has resulted in the part, as it is cooling, shearing glass shards from the plate or breaking it into multiple pieces. In addition to the raw material cost of Victrex AM^TM^ 200 filaments, an almost certain chance of destroying the build plate on every print became cost-prohibitive for an iterative design approach. Vision Miner’s carbon fiber build plate, matrix bonded using aerospace-grade resins, provides a stable foundation that will not break, chip, or shatter operating at ambient temperatures up to 200 °C required for high-performance thermoplastics, remaining compatible with nozzle extrusion temperatures exceeding 450 °C. The build plate is available at approximately 60% of the cost of a 1 kg spool of Victrex AM™ 200 filament. Controlled ramp-down from the ∼100 °C chamber temperature to room temperature over several hours minimizes warping from uneven cooling while differential volumetric shrinkage between the part and build plate naturally shears the two apart without damage or manual intervention.

It was found slight over-extrusion (102% flow) combined with an elevated printing temperature of 440 °C at 100% infill density produces optimal mechanical performance and increased interlayer adhesion in Victrex AM^TM^ 200, at a marginal cost to skin and surface quality. The left 0.6 nozzle of the dual extrusion Funmat Pro 410 was used for the flat thicker base section of the cutting guide to greatly speed up the printing time. The right 0.25 mm nozzle was used for the thin, tall columns and allowed for higher precision in the XY plane and column separation. The first layer with the 0.25 mm nozzle (0.15 mm layer height) overlapped entirely with the last printed layer of the 0.6 mm nozzle (0.25 mm layer height) with reduced flow (85%) and the same hot printing temperature (440 °C). This transition layer is critical to the structural integrity of the column by embedding the first column layer into the last 0.6 mm nozzle surface, the elevated printing temperature locally remelts the surrounding base material, promoting polymer chain interdiffusion across the interface achieving maximum interlayer adhesion at what would otherwise be the weakest interface of the entire part. The approach is analogous to surface ironing concurrently with material deposition. With 4 mm diameter columns, 82.5 mm long and 0.6mm gaps between, default print settings were resulting in oozing/stringing consistently enough to connect all the columns together, leaving no room for the pathologist’s knife. Nozzle build up from oozing and part contact during travel caused the columns to get laterally displaced while printing leading to dimensional inaccuracies, decreased surface quality and column fracture from nozzle bending.

To ensure there is no stringing between adjacent columns or nozzle build up when printing, coasting was used with a retraction and large z hop for all travel between columns. It was found that the coasting volume needed increased from the recommended, (0.25 mm)^3^ = 0.015 mm^3^ (based on nozzle diameter), to 0.125 mm^3^. The resultant minimum volume before coasting (0.3 mm^3^) must be equal to or larger than the coasting volume, ensuring coasting only applies to the walls and not the infill (8 mm travel at 0.15 mm layer height and 0.25 mm line width). This ensures 100% infill with minimal air gaps in the small diameter columns and no stringing while traveling at the high temperatures this material requires. Iterative parameter optimization found the following changes in slicer settings the most relevant to solid infill, long, thin columns with all parameters held constant across print height: retraction minimum travel was decreased to 0.4 mm, coasting volume increased to 0.125 mm^3^, Z hop height increased to 0.75 mm and speed 30 mm/s, travel speed increased to 350 mm/s, and coasting speed decreased to 20 mm/s. With 2 walls, the wall order of operations for column printing is as follows: 1) infill (center of column first), 2) then outer wall to ensure dimensional accuracy, 3) then inner wall, with decreased flow and retraction before coasting within the model geometry to minimize stringing and 4) z hop from the inside wall of the finished column to the center infill of the adjacent column. The circular pathing of the outer and inner walls after nozzle pathing from inside one column to the adjacent will wipe any stringing from the previous high z hop. Printing temperature for the bottom of the column (initial 8 layers, 1.2 mm) was ramped down from 440 °C to the recommended 380 °C to prioritize interlayer adhesion initially before prioritizing decreases stringing for the remainder of the column. Print speed was also decreased from 60 mm/s with the 0.6mm nozzle to 10 mm/s for all column sections.

While formal benchtop mechanical testing was not conducted, extensive iterative prototyping was performed using a range of 3D printing slicer settings and design modifications. Each prototype underwent visual inspection and manual testing for mechanical robustness, including repeated assembly, disassembly, and deliberate fracture to assess failure modes. Optimization was further guided by a comprehensive review of relevant material’s science and additive manufacturing literature, ensuring alignment with current best practices for producing high-performance MRI-compatible components [44–46, 53].

### Embedding and Imaging Protocol

The human postmortem brains, usually left hemispheres without cerebellum or brain stem, are extracted and fixed in 10% formalin (for ADRC Pittsburgh cases) or 4% paraformaldehyde (the Down syndrome cases outside Pittsburgh) for an average of 3 weeks before imaging. The 3D printed enclosure and cutting guide loosely conforms to the shape and size of the left hemisphere of brains of varying sizes with the cerebellum removed. Agar/sucrose mix is poured into the base and allowed to cool as a thin layer, separating the cutting guide from sealing itself to the base.

The cutting guide is inserted into the base followed by the brain. The brain, prior to embedding, has previously been prepared by injecting heated agar into the deepest grooves and ventricle in an effort to eliminate air bubbles. Cooled agar is poured into the container and around the brain at approximately 45 °C to ensure the agar stays viscous while not damaging the tissue to be studied. The neuropathologist gives experienced attention to the filling process, slowly rotating and slightly deforming the brain within the container to further minimize air bubbles left in the sulci of the brain and ventricles. The lid attaches to the base with hardware, more agar is added until the container is filled, and the screw cap is secured. The container is then vacuum sealed to prevent any fluids leaking from the brain or agar from contacting any coil components. For consistent signal while scanning, the container is cooled slowly for several hours until it has reached ∼room temperature [54].

In this work, the images were scanned at 7T using customized head coil of the Tic-Tac-Toe family (Tac G1 and Tac G2) [55–59], which are designed for homogenous and load-insensitive imaging performance in-vivo. The container gets positioned as deep as possible into the head coil; note that it is rotated clockwise slightly (See Supp Fig 4) for improved *B*^+^in the Tac family coils. The size of the container is also compatible with commercial head coils for 7T or lower field strengths. The MRI sequences were optimized for optimal signal at room temperature. The first acquisition is a *B*^+^map (See Supp Fig 4) using a Turbo-FLASH sequence with TR/TE = 2000/1.16 ms; TA = 12 min; flip angle from 0° to 90° in 18° increments; and 3.2 mm isotropic resolution. This 12 minute *B*^+^ map is used to define the optimal reference voltage for the subsequent scans. Next is a 32 minute MP2RAGE with TR/TE = 6000/4.1 ms; Flip Angle = 6 and 7 degrees; TI1 = 514; TI2 = 2020 ms; and 0.4mm isotropic resolution. A 37 minute GRE is then acquired with TR = 40ms; TE1 = 8 ms; TE2 = 15 ms; TE3 = 21 ms; and 0.37 mm isotropic resolution. A 46.5 minute T2-SPACE is acquired with TR/TE = 3400/363 ms; and 0.36 mm isotropic resolution. Diffusion MRI was acquired using the SSFP sequence [7, 60] with the following parameters: TE/TR 21/29 ms, flip angle 24, resolution 1.5 mm isotropic, 2 shells at q-values 150 and 300 cm^−1^, 10 directions each, plus 2 non-directional acquisitions totaling 24 minutes for the DW-SSFP and 156 minutes for the entire protocol.

## Results

### Container Version 1

3 mm columns led to frequent breakage during both printing and use, prompting an increase to 4 mm diameter columns for improved robustness. The vacuum fill port and 3D-printed hose barb adapter were not successful in implementation due to 1) the small hole diameter of the fill port allowed very little agar to be pipetted in before overflowing, and 2) vacuum pressure applied to the hose barb adapter sheared the lid along layer lines before any successful reduction of air bubbles. Sharp corners in the design caused repeated punctures of vacuum seal bags during preparation and imaging, resulting in the potential for fluid leaks. Despite the working features, frequent punctures and mechanical failures highlighted the need for further refinement and material enhancement.

### Dual Cutting Guide Version 1

The axial guide columns presented significant challenges. The short column length fully encased under agar created difficulties in aligning the knife accurately between columns with a 0.5 mm knife and only a 0.55 mm gap, often leading to deformation of the agar and brain or requiring multiple attempts to achieve a clean cut. However, the presence of additional columns necessary for axial cutting reduced the available brain volume in the AP direction by approximately 12.5 mm, significantly limiting the usable internal space of the container leading to the exclusion of several larger brains from this imaging protocol.

### Container Version 2

The increase in gap between columns from 0.55 to 0.6 mm, in addition to increasing column diameter from 3mm to 4mm, was pivotal in extending the operable life of the cutting guide. With increased freedom for positioning the container, optimal positioning to achieve the most homogeneous *B*^+^field was now another variable to consider and was found a slight ∼5-10° tilt (clockwise when inserted into the coil) increased homogeneity. The use of PC, although an improvement from ABS, still led to unwanted deformation of the lid under screw tension leading to a poor seal, leaking agar and heat set inserts pulling loose after dozens of uses. V2 and V3 prototypes transitioned to PC, cutting guide columns still proved highly susceptible to interlayer delamination and column fracture occurred during both fabrication and repeated clinical use. PC, which offered improved mechanical strength and MRI compatibility over ABS, remained prone to low-cycle fatigue failure with relatively low stress, chemical degradation under repeated formalin, bleach and ethanol exposure. Progressive buildup of an unidentified material was observed, likely due to inadequate cleaning between uses and FFF material porosity. Rounded corners eliminated vacuum seal bag puncturing that occurred using container V1.

### Dual Cutting Guide Version 2

All improvements from container version 2 implemented with the dual cutting guide made it far more usable and lasted longer. However, the previously listed limitations, along with the diminished quality of the coronal slabs after replacing the axial slab (See Supp Fig 5), led to the abandonment of the dual guide in favor of higher quality coronal slabs for greater simplicity, reliability, and maximized internal volume. Although this design allowed for nearly 360 degrees of single slab cutting orientation enabling versatile sectioning approaches, the simplicity, robustness, and ease of use of the single cutting guide exposed that the limitations are so fundamental to this design that further development could not overcome them.

### Container Version 3

Despite notable improvements in the changes made in V3, the more complex two-piece snap fit lid leaked and proved unwieldy. The embedding process became more cumbersome, with increased risk of trapping air bubbles, poor brain placement, less secure snap fit causing fluid leakage, directly due to the necessity of two step agar pouring and the more complex lid geometry. The cutting guide’s slanted outer posts and a more rounded base geometry were beneficial and were subsequently incorporated into the next version. The minimal advantages in artifact reduction and field homogeneity of V3 container hardware elimination came at the cost of increased operational complexity for the neuropathologist and a higher propensity for user error during embedding, as well as snap fit lid failure when handling/scanning. However, the ellipsoid external shape of V3 allowed the device to be positioned deeper within the coil for increased transmit field intensity and receive coil performance. An animation comparing V1, V2, and V3 is included in the supplementary information (See Supp Fig 1).

### Container Version 4

The final column diameter are 20 microns smaller than the 4 mm CAD design. The high strength, stiffness, and creep resistance of the Victrex enables the seven screws to be securely tightened, ensuring a leak-proof seal, while also preventing heat-set threaded inserts from dislodging even in the very thin-walled regions. The Victrex cutting guide has no measurable degradation or wear from repeated use cycles and bleach sterilization and an operationally indefinite product life cycle under anticipated conditions of use. High imaging performance was obtained with a *B*_1_^+^ coefficient of variation (CV) of ∼0.259 and average *B*_1_^+^ of ∼624.5 (flip angle/500V) (See Supp Fig 4), calculated averaging two cases using container V4.

### Imaging

Representative images of MP2RAGE, SPACE, GRE, and DW-SSFP are presented **(Fig 7)**. The knife cuts between columns, in either the axial or coronal orientation (See Supp Fig 5). When photographed after cutting, the locations of the cuts are faintly visible and used as the references for MRI registration.

**Figure 7.**
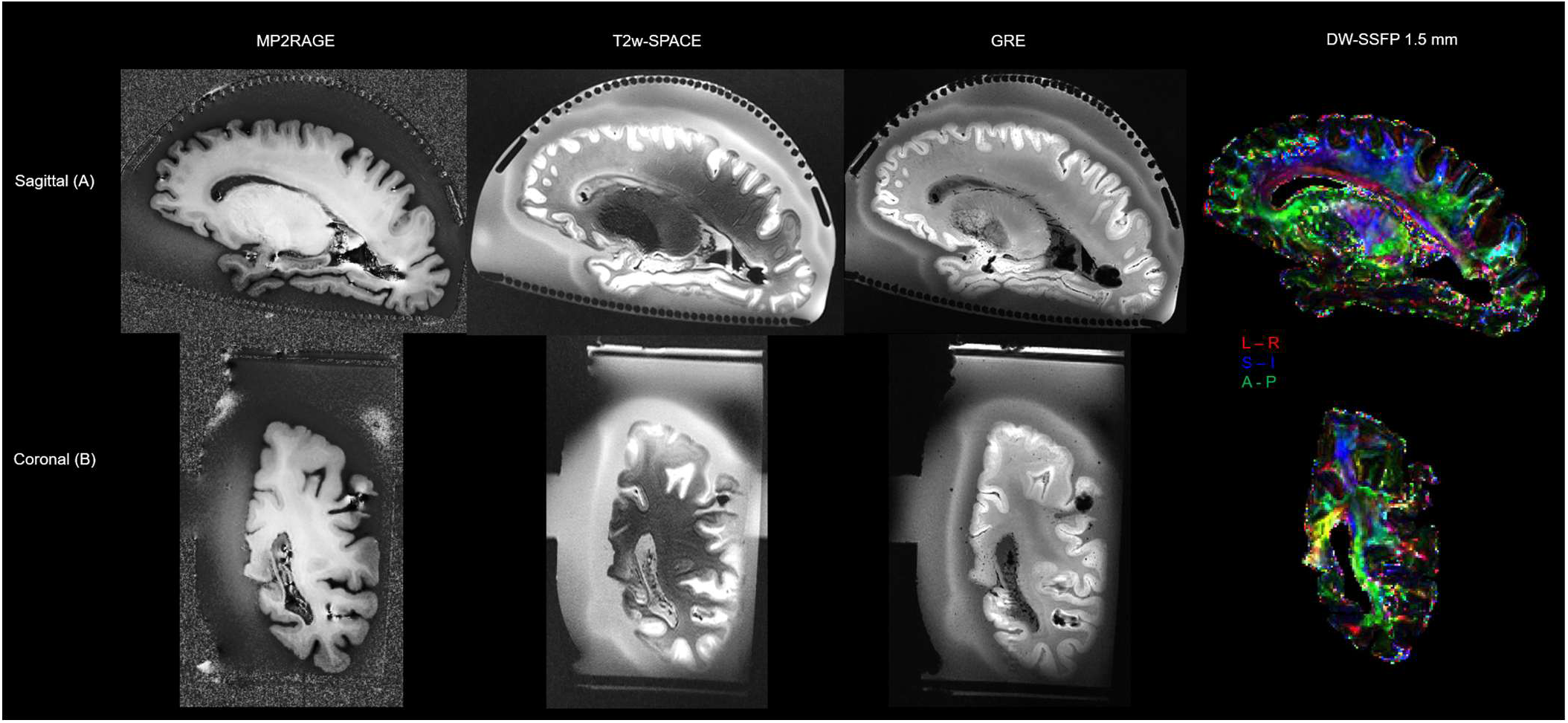
Representative MR images acquired using Container Version 4. Raw images of T1w MP2RAGE, T2w SPACE, T2* GRE, and DW-SSFP for visualization and quantification of brain morphometrics, white matter hyperintensities, cerebral microbleeds, and white matter microstructural changes in (A) sagittal and (B) coronal views.

Cutting between every column will result in a slab thickness of 4.3±0.6 mm. Our current protocol has our neuropathologist cutting between every other column for a slab thickness of 9.2±0.6 mm (Fig 8). After deciding which two columns and the angle to cut, the cutting guide provides the pathologist with the freedom to start at any location with a variety of angles through a singular location. Cuts are made throughout the brain, and a photo of the top surface is taken before any of the slabs removed. The cuts are visible in the image and can be used to identify which columns the knife cut between. The slabs are then removed and individually imaged for block face photographs of the gross anatomy. The MR images are then able to be oriented to re-slicing in the coronal plane perpendicular to the flat base of the cutting guide. The marks on the surface of the agar are visible (**Fig 8A)** and indicate which two columns on the top and bottom row are used to cut between, and the image is resliced as though the knife traveled from center of the gap between columns on the top and bottom rows. Figure 8B shows the alignment of the MR images with block face gross anatomy photographs to sample WMHs for histological sampling.

**Figure 8.**
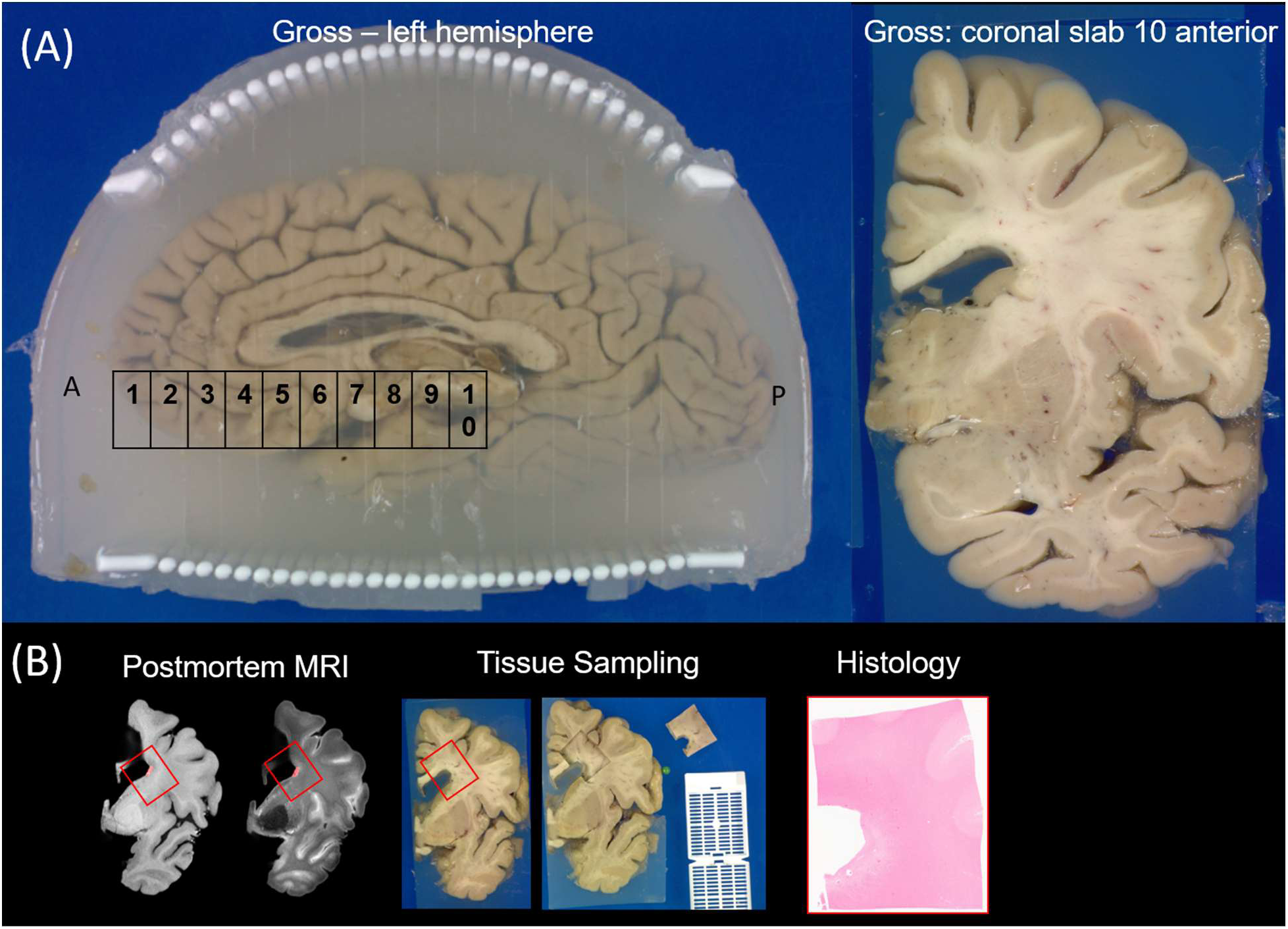
Container Version 2 (pictured) enables postmortem brain tissue cutting, targeted tissue sampling, and subsequent histology. (A) The left hemispheres are cut coronally, and the image represents the anterior surface of the coronal tissue slab number 10, counting from the anterior side of the brain. (B) With MR and tissue slab images, co-registration occurs using Rembg, rigid rotation and scaling to co-register postmortem MRI and tissue slabs for targeted tissue sampling and subsequent histology.

Container V4 is now implemented in ongoing research and brain bank protocols, providing a reliable platform for postmortem MRI-histology correlation at 7T for 215 human brains **(Table 2)**.

**Table 2.**
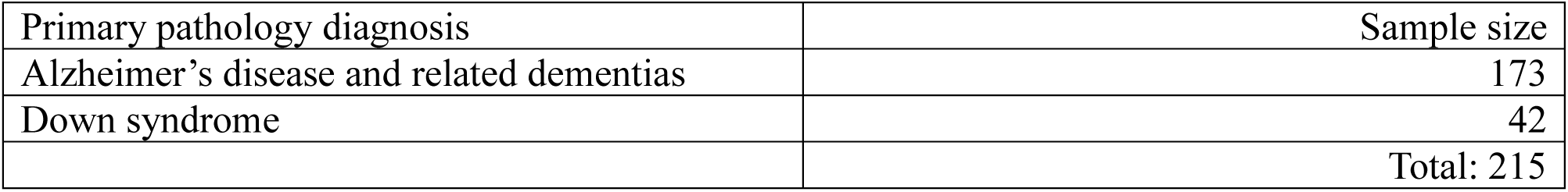
Breakdown of 215 postmortem brains imaged for MRI-histology correlation as of April 30, 2026. Most postmortem brains have usable MR images of T1w MP2RAGE, T2w SPACE, and T2* GRE; only the most recent 46 postmortem brains have MR images of DW-SSFP.

| Primary pathology diagnosis | Sample size |
| --- | --- |
| Alzheimer's disease and related dementias | 173 |
| Down syndrome | 42 |
|  | Total: 215 |

## Discussion

Recent advancements in digital neuropathology and image analysis, as shown by Julian et al. [34] and many others [61, 62], have highlighted the growing importance of large-scale, spatially accurate, and histologically validated datasets for neurodegenerative disease research. However, a limitation in this field remains the ability to spatially correlate microscopic pathology with imaging markers consistently [34, 62]. The stereotaxic enclosure and cutting guide system presented in this work directly addresses this gap by enabling more precise MRI-informed sampling of postmortem brain tissue with minimal deformation. By preserving the brain’s orientation from high-resolution 7T MRI through to histological sectioning, this platform facilitates accurate ground-truth labeling of radiographically visible lesions such as WMHs, microinfarcts, and quantifications of cortical thickness and regional volumes in the hippocampus and the amygdala furthering Alzheimer’s disease and related dementias research [63]. The container design and embedding protocol was proved successful, with 215 brains scanned to data as part of AD and DS studies, which is corroborated by additional studies and publications utilizing the container [14, 32–35, 64, 65]. Additionally, electromagnetic properties of average *B*^+^field and CV, essential for high quality imaging, are found to be comparable to industry standard in-vivo 7T transmit coil performance (See Supp Fig 5).

The stereotaxic brain enclosure and cutting guide described in this study directly enabled the high-fidelity MRI-histology correlation achieved in recent neuropathology case reports investigating cognitive resilience in an individual with Down syndrome despite significant Alzheimer’s disease pathology [64]. As shown in the study, spatial fidelity was essential for validating imaging signatures against detailed immunohistochemical findings, contributing to insights on brain reserve and resilience mechanisms.

Another major benefit our method provides unlike other protocols is that the brain is scanned only once intact with minimal deformations using agar as a medium to suspend the tissues fixed in place in relation to the cutting guide. This is the closest feasible ex-vivo replication of in-situ conditions for high throughput data collection. The most prominent research in guided cutting of brain sections for MRI registered histological examinations requires two instances of scanning and manufacturing of a custom shaped cutting guide based on the structural information for the first MRI session [17, 19, 20, 24, 28, 30, 66, 67].

This device’s potential extends beyond current applications, offering significant insights into other brain phenomena that are indistinguishable in gross anatomy. For instance, cortical microinfarcts, which are small, often subclinical infarcts, can be visualized on high-resolution MRI scans as areas of cortical thinning or signal changes. Histological examination of these microinfarcts can identify microvascular pathology, neuronal loss, and gliosis, thereby enhancing our understanding of their impact on cognitive decline and neurodegenerative diseases.

The deep sulci of the brain and ventricle cavity pose a unique challenge not all organs are subjected to. The device’s principal functions can be applied to other non-chambered organs, such as the liver, spleen, pancreas, and kidneys, to study many phenomena distinguishable by MRI but not by gross anatomy. Beyond the brain, this device supports MRI-histology correlation in other organs where pathology may be missed on gross inspection. In the liver, MRI can detect steatosis and fibrosis, while histology confirms cellular changes and progression [68]. The pancreas benefits, similarly, with early fibrosis and chronic pancreatitis visible on MRI but requiring histology for detailed characterization [69]. In the kidneys, cortical microcysts and fibrotic changes appear on MRI as subtle textural alterations, with histology aiding early diagnosis [70]. MRI permits characterization of a wide range of lesions and vascular abnormalities, including infarction, cysts, hemangiomas, and other pathological changes amongst many organs such as the spleen, with imaging features that reflect underlying tissue pathology [71]. These applications extend the device’s utility across multiple disease models and organ systems.

This study has limitations. The current procedure has challenges in repeatably eliminating air bubbles with 7T brain MR imaging using semi-solid gel agar but this approach still offers greater specimen stability and fewer global imaging artifacts than fluid-based immersion protocols, which still grabble with difficulties in regards to air entrapment in addition to motion artifacts [72]. Air bubbles create artifacts that impede comprehensive grey matter MRI studies. Agar bonds to a small portion of the outermost layer of grey matter, further compromising histological studies of the outer-most cells of the brain. The brain is carefully massaged during the introduction of the agar in its semi-solid state, thereby mitigating this issue, although not eliminating it completely. It is worth noting that this does not impede the ability to identify, segment and register WMHs to gross anatomy [14]. Moreover, Ex vivo imaging involving, loss of cranial pressure, paraformaldehyde fixation, tissue hardening/dehydration/rehydration in fixation all contribute to a postmortem tissue deformation and signal changes [73] that can make partial and full brain volume analysis inaccurate when compared to in vivo or in situ [27, 74] but methods such as non-linear distortion correction methods using prior in vivo scans when available to improve accuracy and reliability of MR image analysis [14].

Future studies in white matter hyperintensity (WMH) lesion segmentation will be conducted leveraging deep learning models and registered to the block face photographs for high special confidence during histological sampling. This process provides a one-to-one correspondence between the MR image and histology free from deformation overcoming the long-standing challenges of registration for these modalities. The robustness of the design and methodology could be further validated with the development of a fully automated pipeline of the current WMHs segmentation and registration to gross anatomy block face photographs would allow for the pathologist to continue to histological sampling immediately after they finish the cutting process. A fully automated pipeline that provides neuropathologists with multiple image sets based on theoretical cutting locations before any sectioning takes place, ensuring the most informed and desirable cuts possible are made. The container design can always be further tailored for specific use cases and scalable design practice. The dual cutting guide could prove useful especially for other organs or when studying alternative brain phenomena. A more robust and user-friendly snap fit lid could be developed through costly continual iterative designs and more accurate mechanical tolerances with the use of Victrex AM^TM^ 200.

## Conclusion

The methods developed through iterating and improving the design of the novel container and cutting guide in conjunction with the use of in-house developed RF coils have led to an extensive brain bank of postmortem brain specimens with histological analysis in addition to retrospective MR studies. The ability to conduct tissue specific histology on regions indistinguishable in gross anatomy with complete confidence enhances our ability to study nature and extent WMHs play a role in neurodegenerative diseases. The novel integral cutting guide and container, acting as a stereotaxic device, not only streamlines the imaging and sectioning process for histological examination of the brain but also holds significant potential for application to other organs and conditions, paving the way for future advancements in medical research and diagnostics.

## Availability of Data and Materials

CAD models of all container versions available at https://github.com/JBelli/Cutting-Guide. Additional data collected supporting the conclusions of this article is available upon request to the corresponding author.

## Abbreviations

ABS: Acrylonitrile Butadiene Styrene
ADRC: Alzheimer’s Disease Research Center
ADRD: Alzheimer’s Disease and Related Dementias
CV: Coefficient of Variation
DS: Down syndrome
FDM: Fused Deposition Modeling
FFF: Fused Filament Fabrication
MRI: Magnetic Resonance Imaging
PAEK: PolyArylEtherKetone
PC: Polycarbonate
RF: radio frequency
TTT: Tic Tac Toe
UHF: Ultra High Field
WMHs: White Matter Hyperintensities
7T: 7 Tesla

## Acknowledgements

This work was supported by the National Institutes of Health under award numbers R01MH111265, T32MH119168, R01AG063525, U19AG068054 and BrightFocus Foundation A2025009F. This research was supported in part by the University of Pittsburgh Center for Research Computing and Data, RRID:SCR_022735, through the resources provided. Specifically, this work used the HTC cluster, which is supported by NIH award number S10OD028483.

## Ethics Declaration

This study was conducted in accordance with the Declaration of Helsinki and all neuropathology study exempt of IRB.

## Supplemental Figures

**Figure S1.**
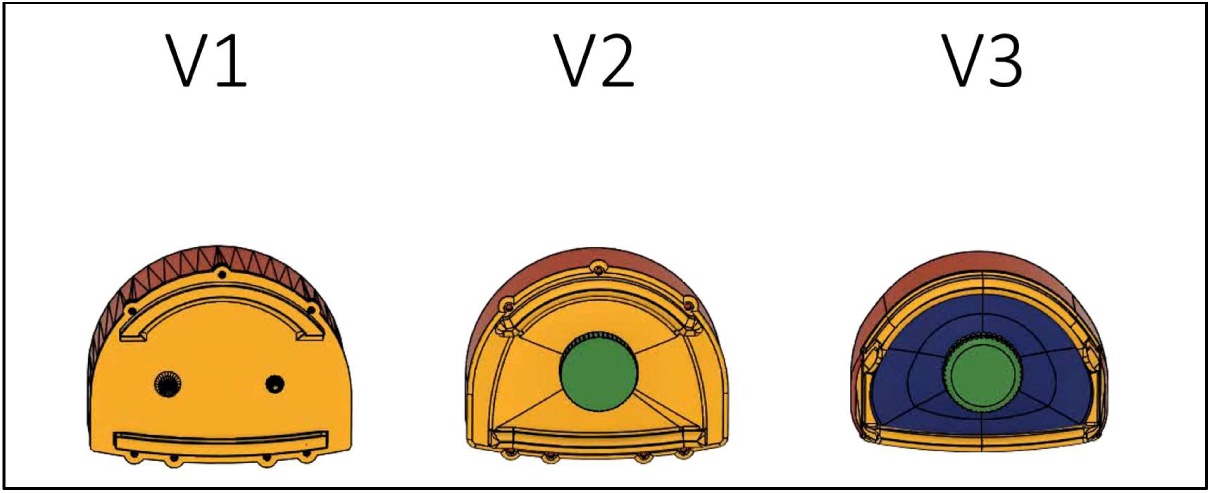
V1 V2 V3 comparison video

**Figure S2.**
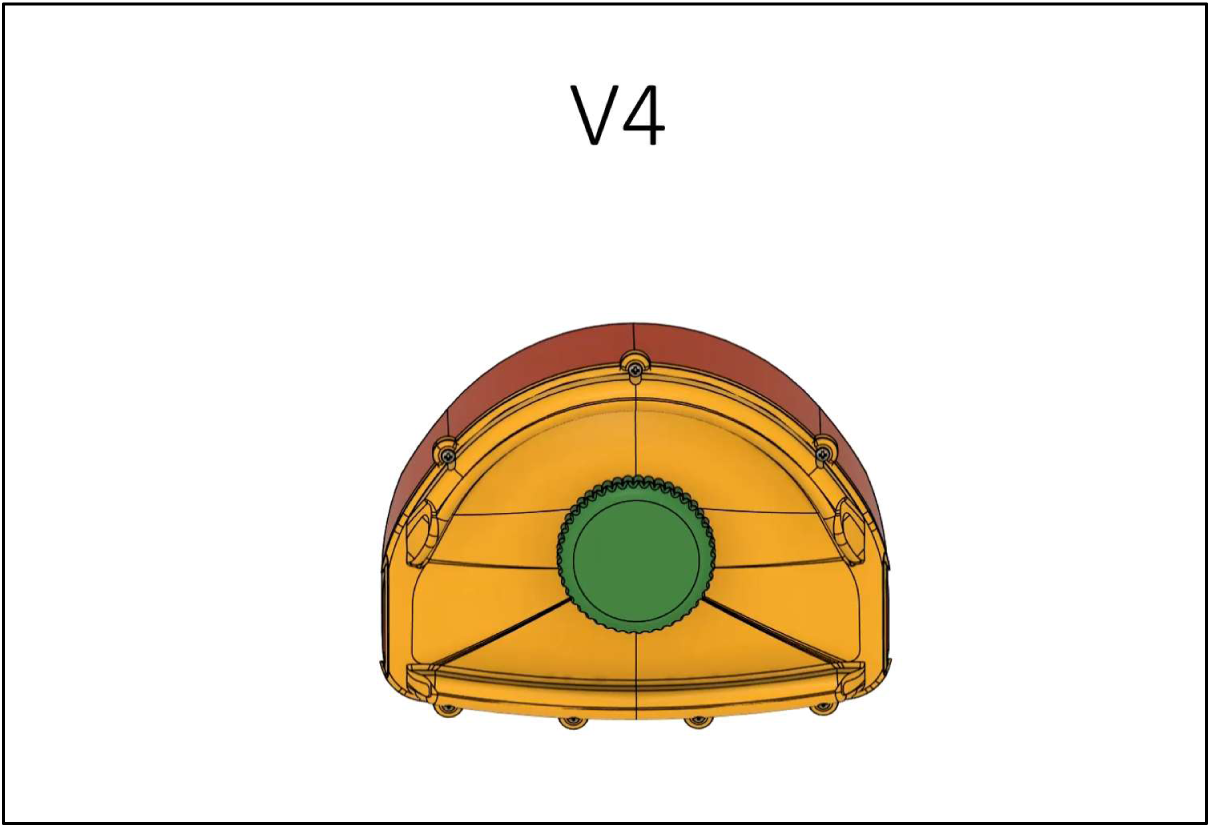
V4 animation video

**Figure S3.**
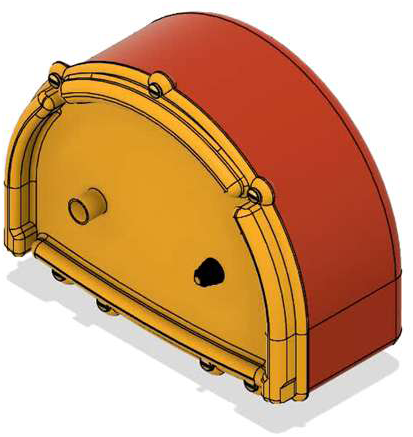
V1 lid V2 base container and dual cutting guide design

**Figure S4.**
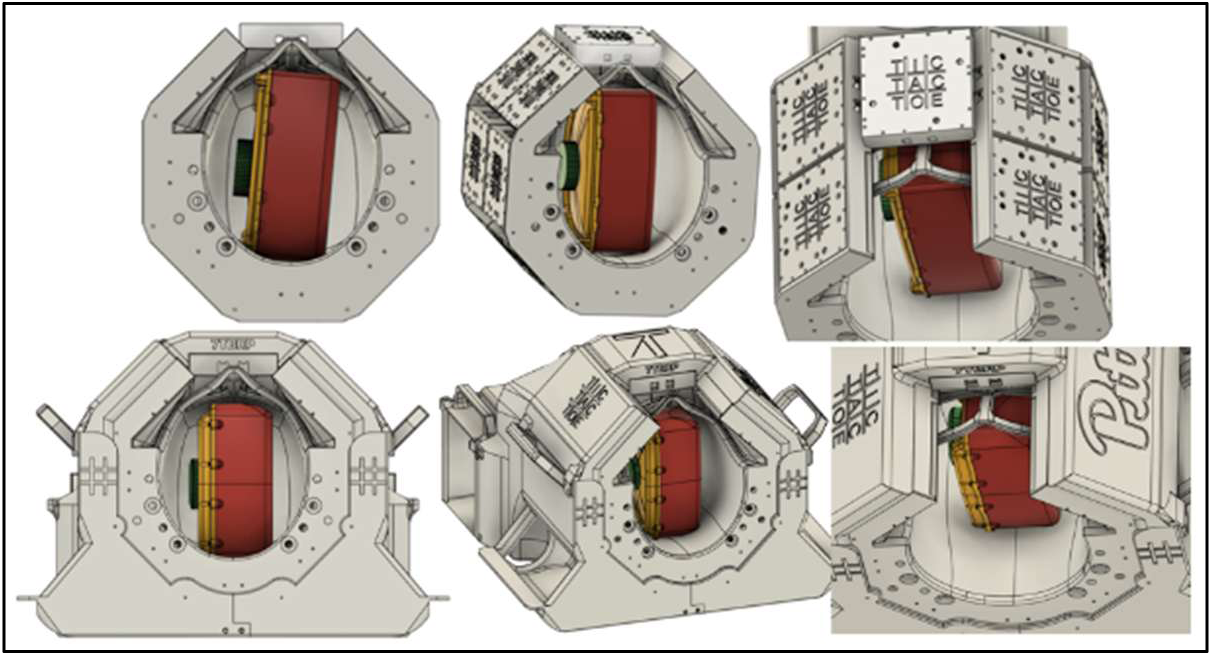
Comparison of Container Version 2 (top) and Container Version 4 (bottom) allow for deeper placement of container in the receive helmet of the head coil.

**Figure S5.**
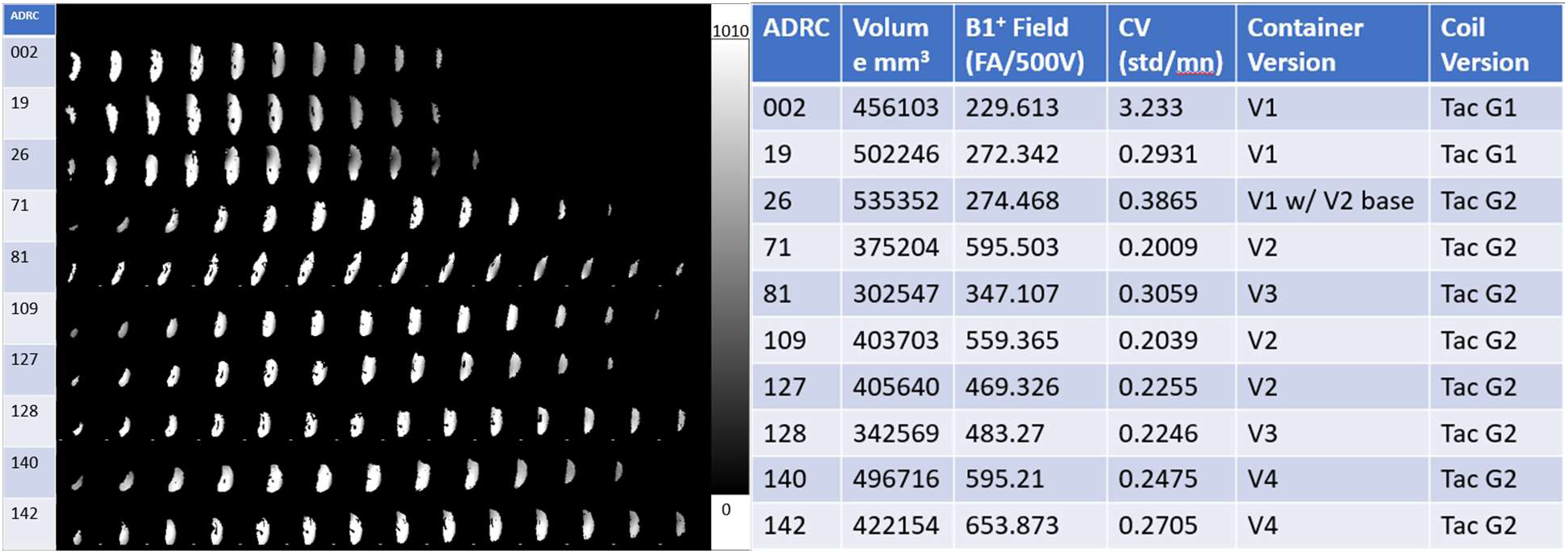
Evolution of the *B*^+^ field distribution across container versions. Beginning April 2021, Case #1 represents first initial prototype of container V1 the Tac G1 (16ch transmit) coil. Case #2 was a brain scanned under the same conditions after the magnet underwent a gradient coil upgrade. Case #3 was done with container V1 lid and cutting guide, and V2 base in the Tac G2 (60ch transmit) coil. Case #4 was processed with the V2 lid, cutting guide and base and a second improved shim case. Case #5 was scanned and cut using the V3 container. Case #6 was scanned with container V2 and a third improved shim case. Case #7 was scanned with container V2, a fourth improved shim case and additional gradient coil upgrade. Case #8 was scanned with container V3. Case #9 and #10 were scanned with container V4 and after a scanner upgrade to the 7T plus system. These were acquired from a *B*^+^mapping sequence and masked using structural scans to represent the magnetic field distribution within the tissue only. The selection of cases represents a timeline from the initial prototype of container V1 to our most recent container, head coil, and 7T scanner. Improved *B*^+^ Field and approximate CV of 0.26 can be seen as comparable for the industry standard 7T in-vivo head coils [41, 75].

**Figure S6.**
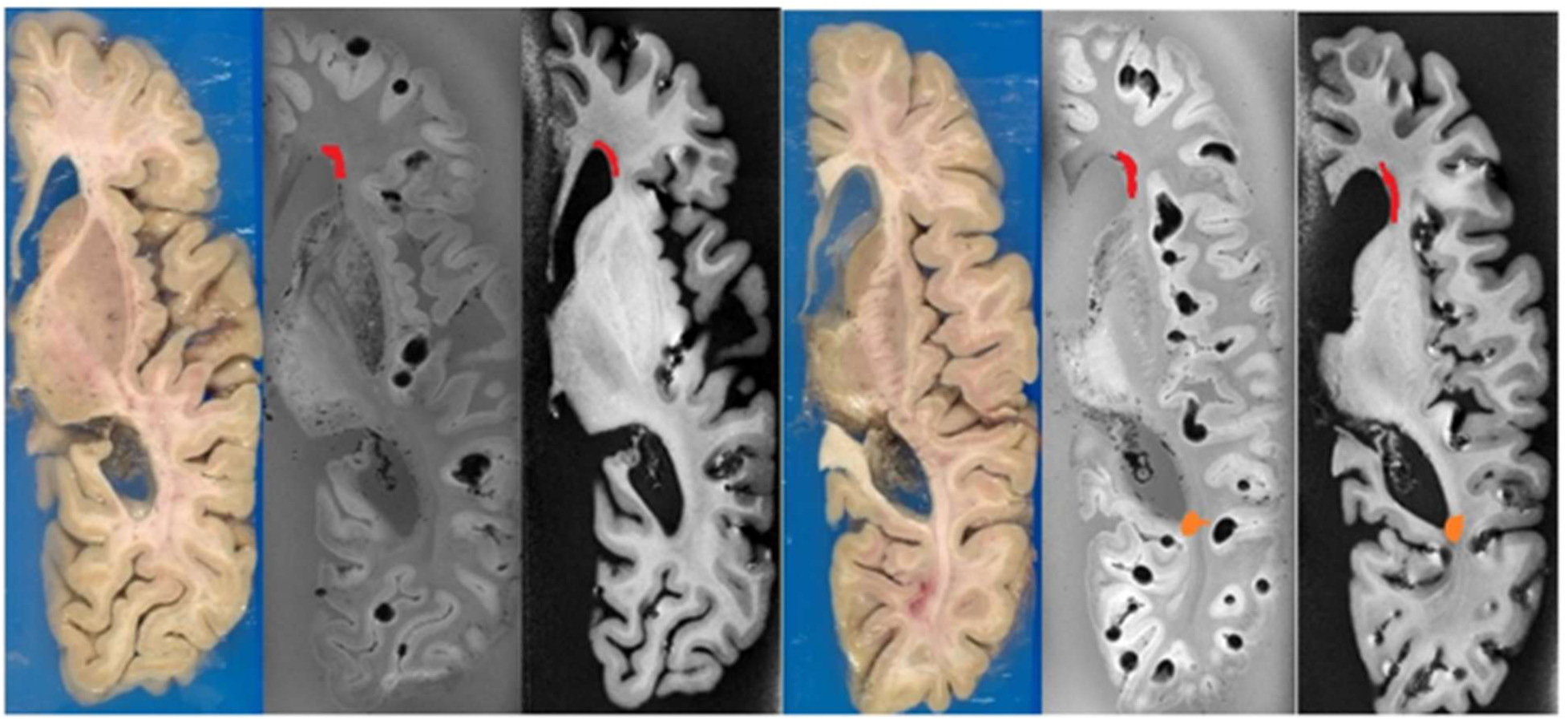
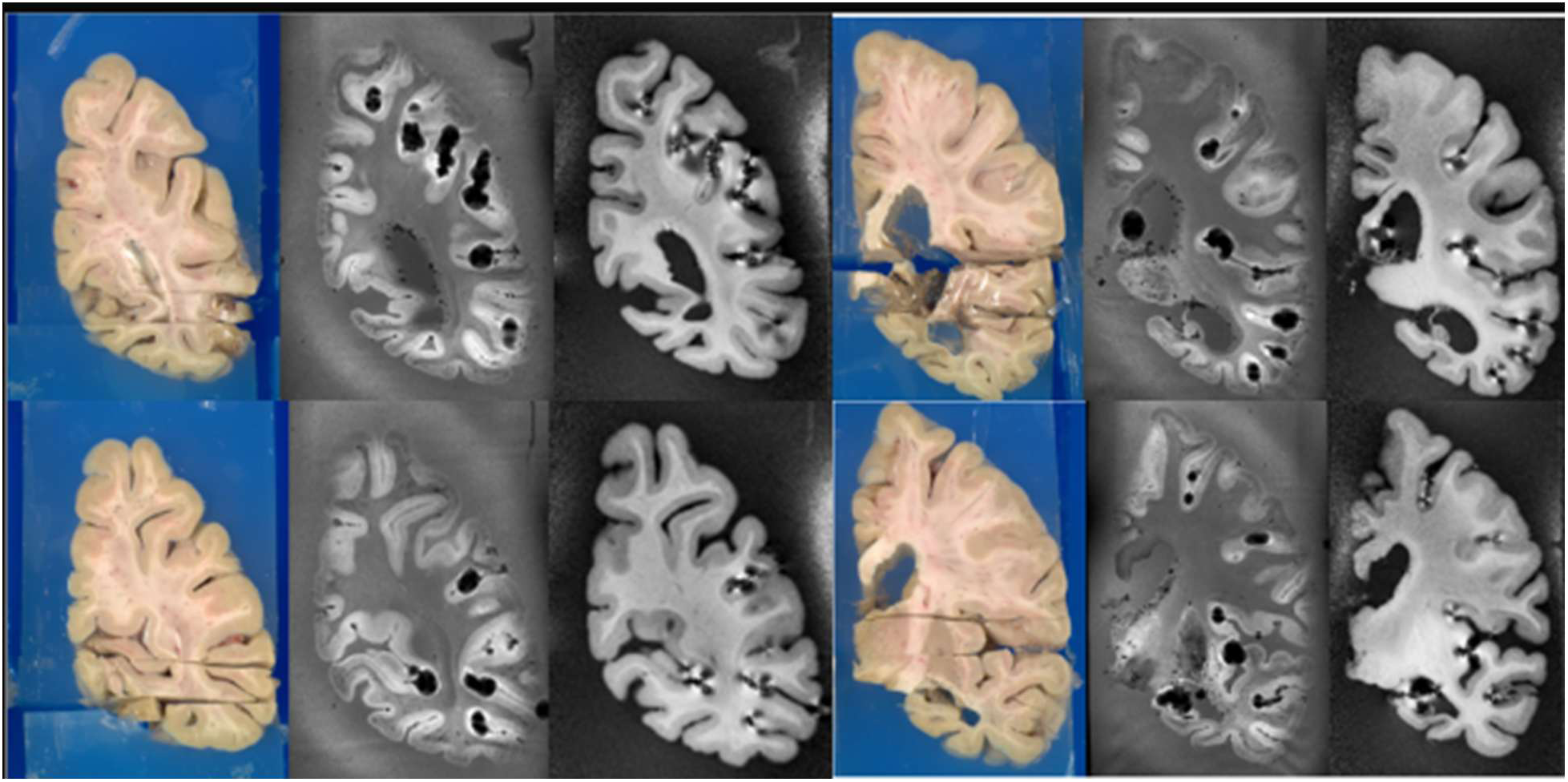
An example of two different brains (top and bottom) with an axial slab removed and WMHs manually denoted in the registered MR images in orange and red. The alignment of the coronal cuts after replacing the axial slab are shown. The axial cuts previously made are visible in the gross anatomy but, due to agar fixation, still align with the MR images.

**Figure S7.**
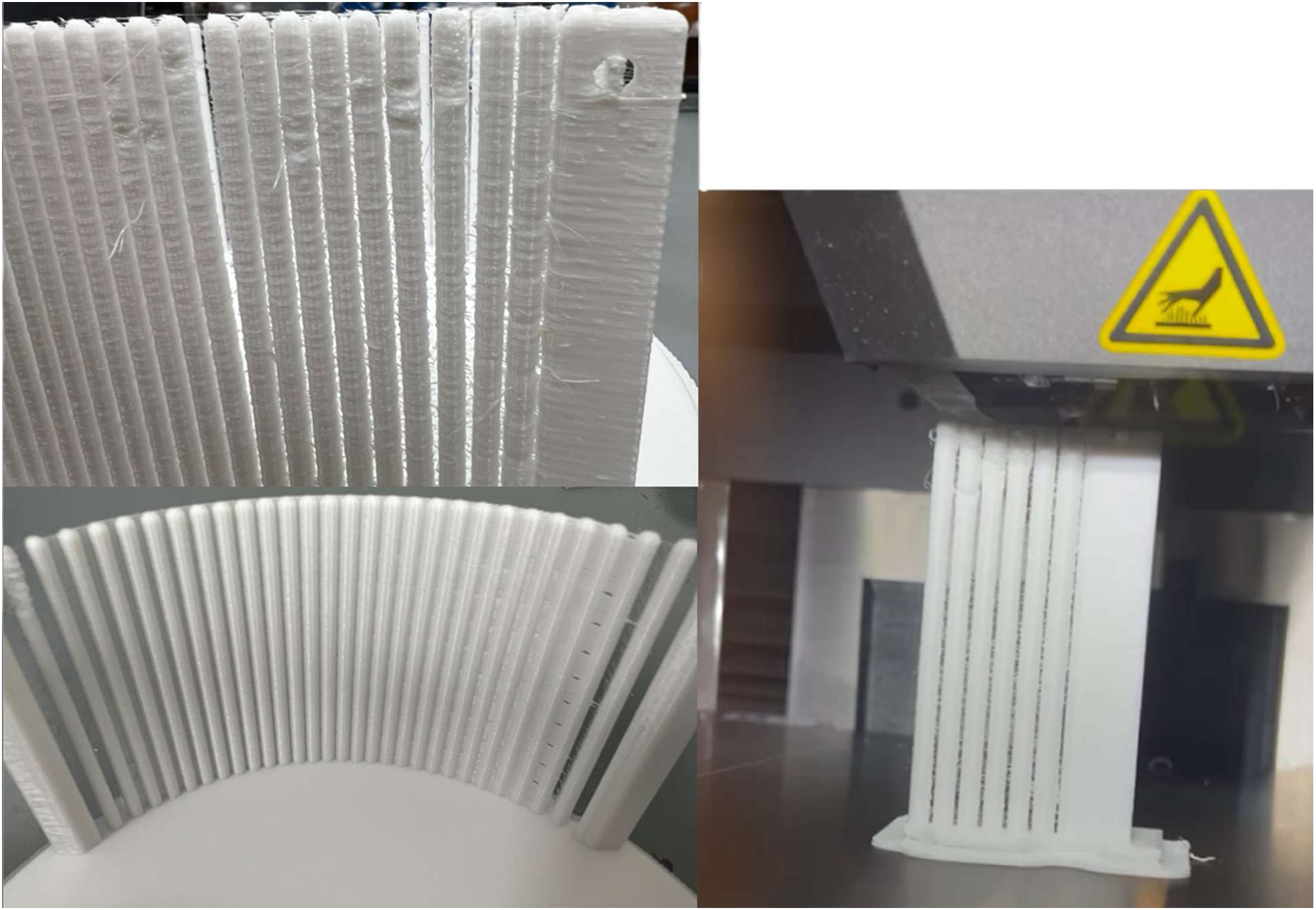
FFF manufacturing challenges associated with tall, thin bending columns. Representative challenges included maintaining dimensional stability of slender column geometries, minimizing stringing between adjacent columns, and reducing ribbing, gaps, or other print defects. Poor column printing results were mitigated with the changes in slicer settings (See *Iterative Design leveraging FFF additive manufacturing*).

