## Supplementary figures and images for "An open-source stereotaxic container with an integrated cutting guide for human brain fixation during magnetic resonance imaging and sectioning for histology"

### Supp Fig 4.png

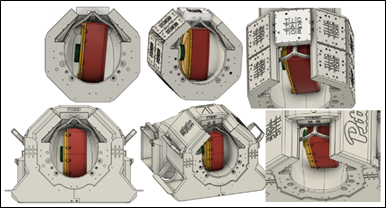

### Supp Fig 5.png

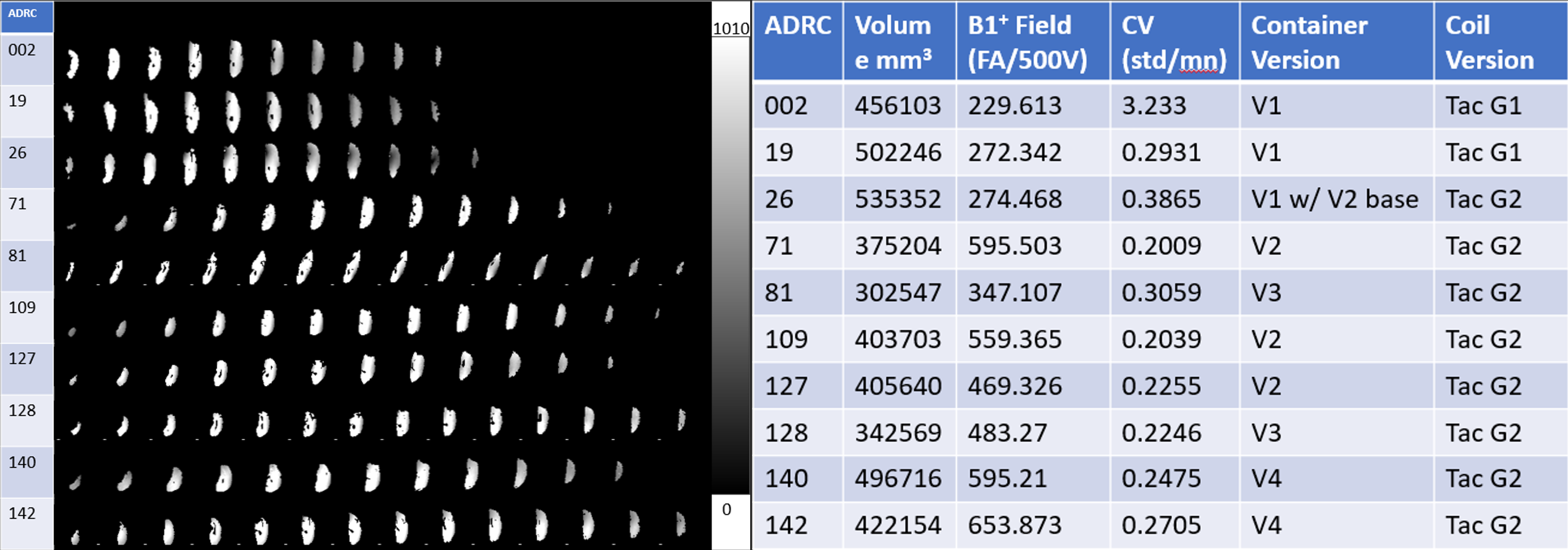

### Supp Fig 6.png

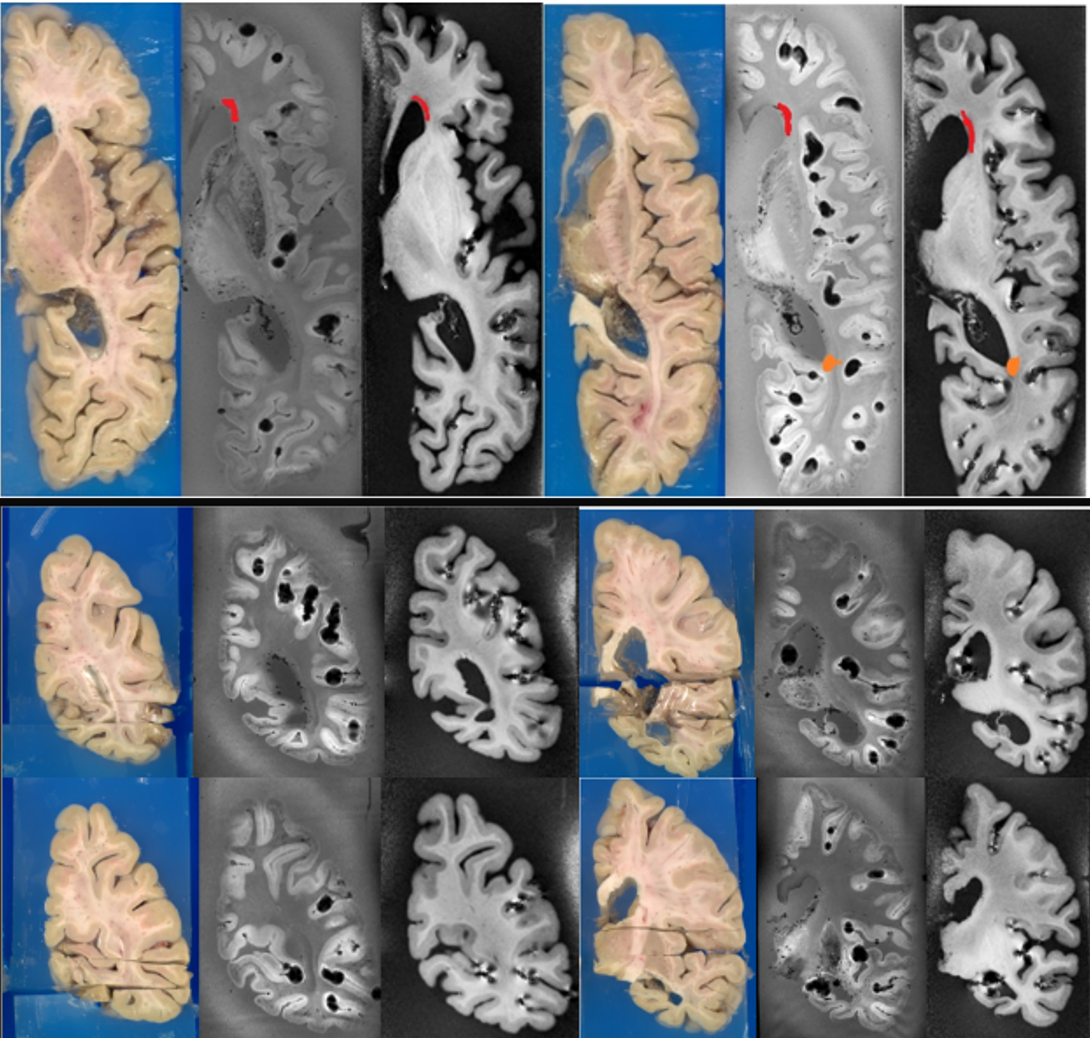

### Supp Fig 7.png

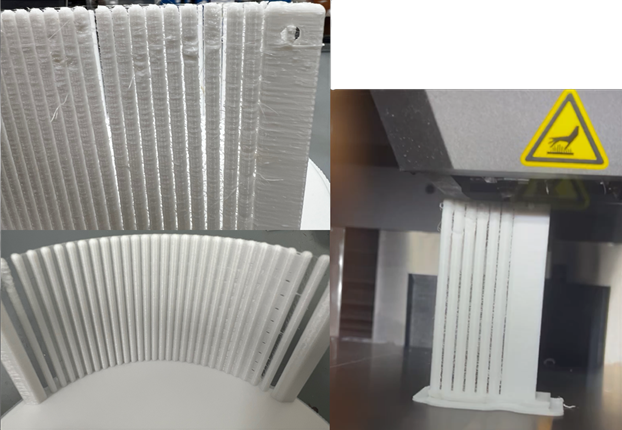
